# Valection: Design Optimization for Validation and Verification Studies

**DOI:** 10.1101/254839

**Authors:** Christopher I. Cooper, Delia Yao, Dorota H. Sendorek, Takafumi N. Yamaguchi, Christine P’ng, Cristian Caloian, Michael Fraser, SMC-DNA Challenge Participants, Kyle Ellrott, Adam A. Margolin, Robert G. Bristow, Joshua M. Stuart, Paul C. Boutros

## Abstract

**Background:** Platform-specific error profiles necessitate confirmatory studies where predictions made on data generated using one technology are additionally verified by processing the same samples on an orthogonal technology. In disciplines that rely heavily on high-throughput data generation, such as genomics, reducing the impact of false positive and false negative rates in results is a top priority. However, verifying all predictions can be costly and redundant, and testing a subset of findings is often used to estimate the true error profile. To determine how to create subsets of predictions for validation that maximize inference of global error profiles, we developed Valection, a software program that implements multiple strategies for the selection of verification candidates.

**Results:** To evaluate these selection strategies, we obtained 261 sets of somatic mutation calls from a single-nucleotide variant caller benchmarking challenge where 21 teams competed on whole-genome sequencing datasets of three computationally-simulated tumours. By using synthetic data, we had complete ground truth of the tumours’ mutations and, therefore, we were able to accurately determine how estimates from the selected subset of verification candidates compared to the complete prediction set. We found that selection strategy performance depends on several verification study characteristics. In particular the verification budget of the experiment (*i.e.* how many candidates can be selected) is shown to influence estimates.

**Conclusions:** The Valection framework is flexible, allowing for the implementation of additional selection algorithms in the future. Its applicability extends to any discipline that relies on experimental verification and will benefit from the optimization of verification candidate selection.

## Background

High-throughput genomics studies often exhibit error profiles that are biased towards certain data characteristics. For example, predictions of single-nucleotide variants (SNVs) from DNA sequencing data have error profiles biased by local sequence context [1-2], mappability of the region [3] and many other factors [4-5]. The false positive rate for individual predictions in high-throughput studies is frequently high [6-7], while the false negative rate is difficult to estimate and rarely known. Critically, error rates can vary significantly between studies because of tissue-specific characteristics, such as DNA quality and sample purity, and differences in data processing pipelines and analytical tools. In cancer studies, variations in normal tissue contamination can further confound genomic and transcriptomic analyses [8-10].

Taken together, these factors have necessitated the wide-spread use of studies with orthogonal technologies, both to verify key hits of interest and to quantify the global error rate of specific pipelines. In contrast to a *validation study,* which typically approaches the same biological question using an independent set of samples (*e.g.* like a test dataset in a machine learning exercise), we define a *verification study* as interrogating the same sample-set with an independent method (*i.e.* a method that generates analogous data using a distinct chemistry). The underlying concept is that if the second technique has separate error profiles from the first, a comparative analysis can readily identify false positives (*e.g.* in inconsistent, low quality calls) and even begin to elucidate the false negative rate (*e.g.* from discordant, high quality calls).

The choice of verification platform is critical as it determines both the tissue and financial resources required. There is typically a wide range of potential verification technologies for any given study. While confirmation of DNA-sequencing results traditionally involves gold-standard Sanger sequencing [11-12], the drawbacks of this approach (*e.g.* high financial and resource costs) and advancements in newer sequencing techniques have shifted the burden of variant verification to other technologies [13-15]. For example, a typical Illumina-based next-generation sequencing (NGS) whole-genome or whole-exome experiment may be verified by sequencing a separate library on a different but similar machine [16]. This offers the advantages of high-throughput, low cost and the opportunity to interrogate inter-library differences [17]. Other groups have applied mass-spectrometric based corroboration of individual variants, which has the benefit of technological independence [18-19].

Apart from choice of technology, all groups must make decisions regarding the *scope* of their verification work. For example when considering genome-wide discovery, it may be appropriate to verify only known candidate drug target mutations or unexpected novel functional aberrations. However, in many contexts having an unbiased estimate of the global error rate is critical. This is particularly true when benchmarking different data-generating methods or when looking at genome-wide trends. It remains unclear how best to select targets for verification studies, particularly in the context of fairly comparing multiple methods and providing unbiased performance metric estimates. To address this problem, we created Valection, a software tool that implements a series of diverse variable selection strategies, thereby providing the first framework for guiding optimal selection of verification candidates. To benchmark different strategies, we exploit data from the ICGC-TCGA DREAM Somatic Mutation Calling Challenge (SMC-DNA), where we have a total of 2,051,714 predictions of somatic SNVs made by 21 teams through 261 analyses [20, 4]. We show that the optimal strategy changes in a predictable way based on characteristics of the verification experiments.

## Results

We began by developing six separate strategies for selecting candidates for verification **(Figure 1).** The first is a naïve approach that samples each mutation with equal probability, independent of whether a mutation is predicted by multiple algorithms or of how many calls a given algorithm has made (‘random rows’). Two simple approaches follow that divide mutations either by recurrence (‘equal per overlap’) or by which algorithm made the call (‘equal per caller’). Finally, we created three approaches that account for both factors: ‘increasing per overlap’ (where the probability of selection increases with call recurrence), ‘decreasing per overlap’ (where the probability of selection decreases with call recurrence) and ‘directed-sampling’ (where the probability of selection increases with call recurrence while ensuring an equal proportion of targets is selected from each caller). All methods have programmatic bindings in four separate open-source languages (C, R, Perl and Python) and are accessible through a systematic API through the Valection software package. Valection thus becomes a test-bed for groups to try new ways of optimizing verification candidate-selection strategies.

**Figure 1:**
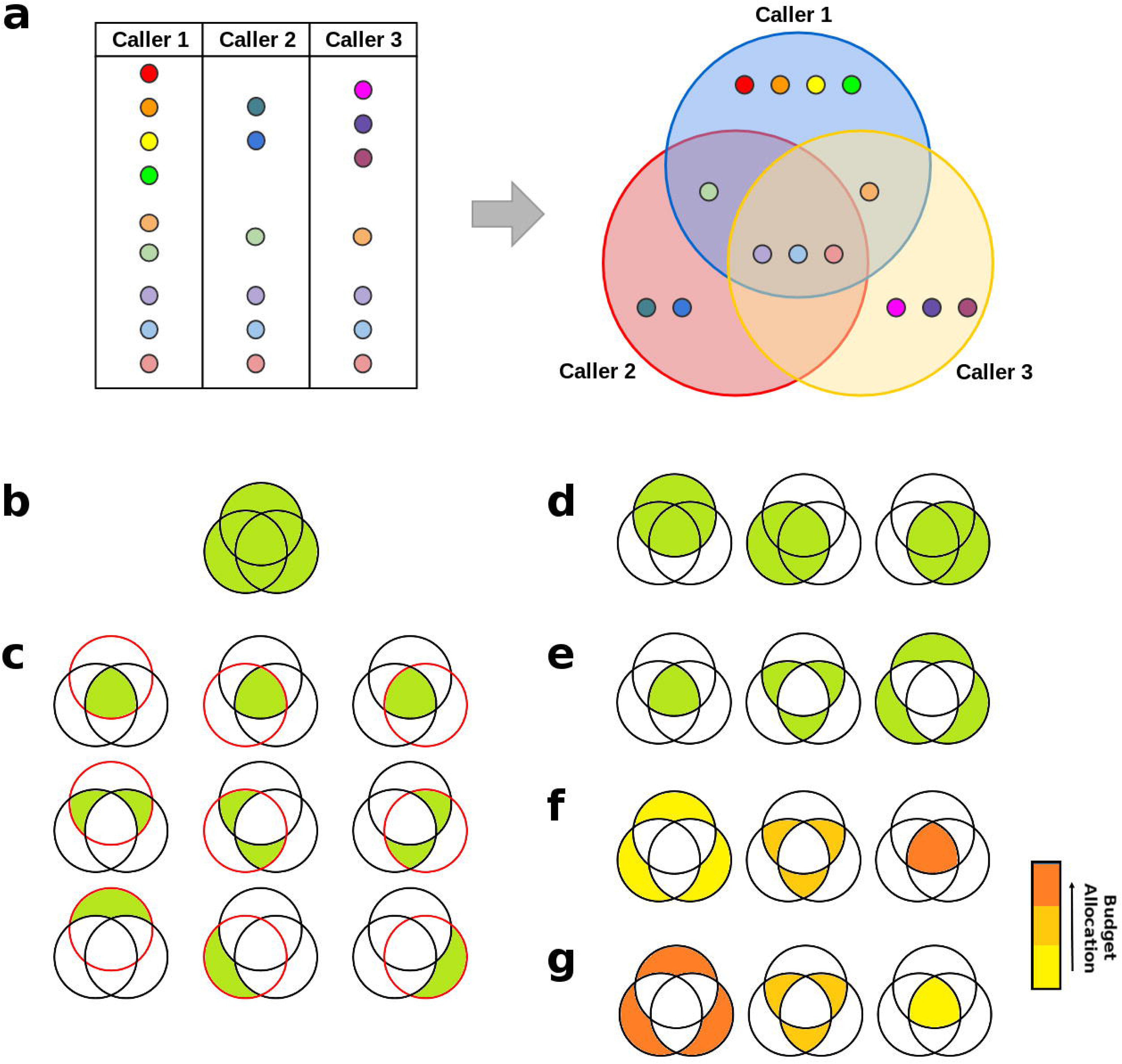
Valection Candidate-Selection Strategies. a) A hypothetical scenario where we have results from three callers available. Each call is represented using a dot. SNV calls that are shared by multiple callers are represented with matching dot colours. b) The ‘random rows’ method where all unique calls across all callers are sampled from with equal probability. c) The ‘directed-sampling’ method where a ‘call overlap-by-caller’ matrix is constructed and the selection budget is distributed equally across all cells. d) The ‘equal per caller’ method where the selection budget is distributed evenly across all callers. e) The ‘equal per overlap’ method where the selection budget is distributed evenly across all levels of overlap (*i.e.* call recurrence across callers). f) The ‘increasing with overlap’ method where the selection budget is distributed across overlap levels in proportion to the level of overlap. g) The ‘decreasing with overlap’ method where the selection budget is distributed across overlap levels in inverse proportion to the level of overlap.

To compare the six methods outlined above, we used data from tumour-normal whole-genome sequencing pairs from the ICGC-TCGA DREAM Somatic Mutation Calling Challenge [20, 4]. These tumours differ in major characteristics such as normal contamination, sub-clonality and mutation rate. We chose to work with simulated tumours because we know the ground truth of their mutational profiles, allowing a precise evaluation of the effectiveness of different selection schemes in estimating the true underlying error rates. Altogether, there are results available from 261 SNV calling analyses performed by 21 teams. We designed a rigorous parameter-sweeping strategy, considering different numbers of SNV calling algorithms and different quantities of verification candidate targets. The experimental design is outlined in **Figure 2.**

**Figure 2:**
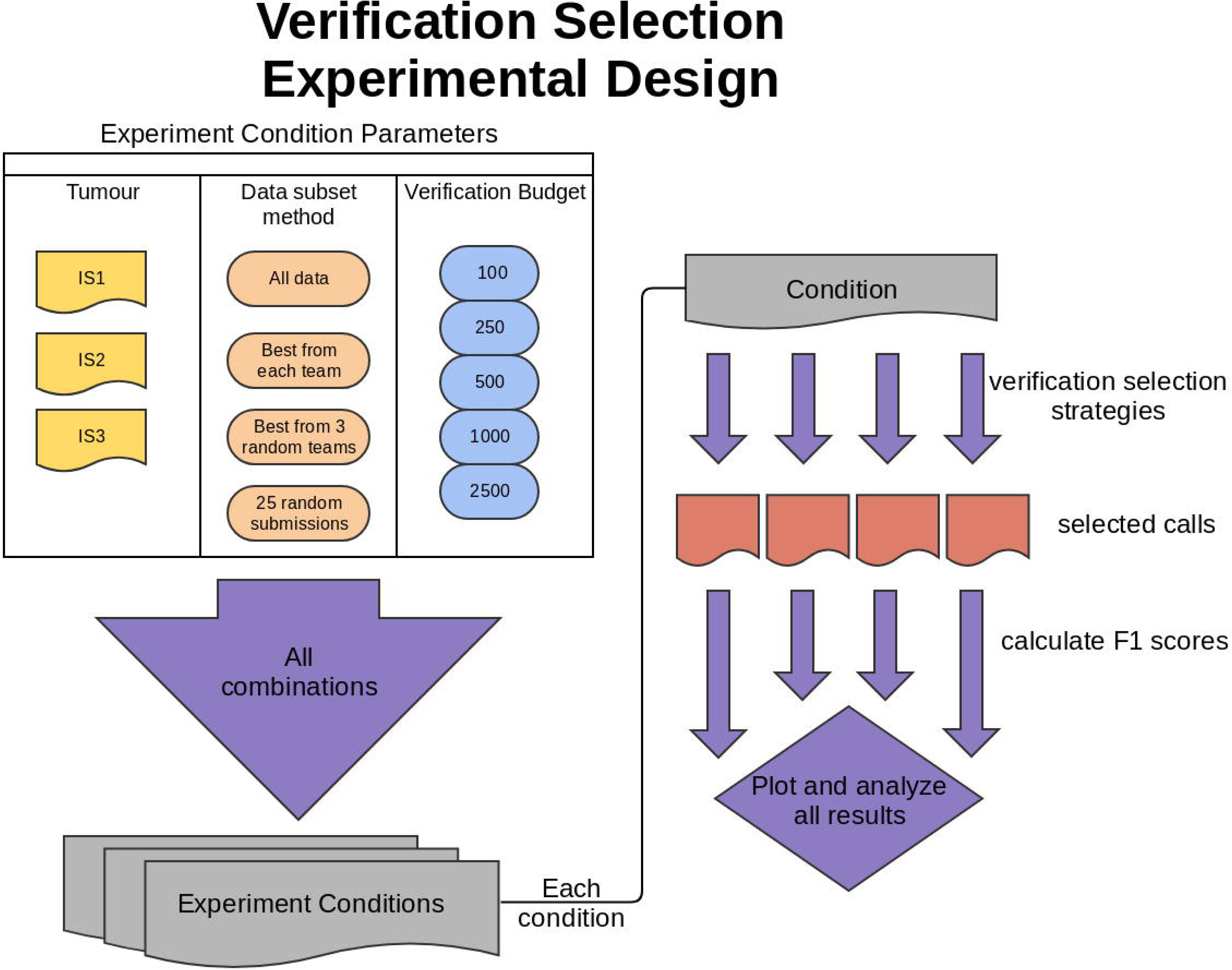
Verification Selection Experimental Design. Verification candidates were selected from somatic mutation calling results of multiple algorithms run on three *in silico* tumours (IS1, IS2, and IS3). Candidate selection was performed separately on each tumour’s set of results using all combinations of five different verification budgets (*i.e.* number of calls selected) and six different selection strategies. F_1_ scores were calculated for each set of selected calls and compared to F_1_ scores calculated from the full prediction set. To compare the effect of the numbers of algorithms used, datasets were further subset using four different metrics.

We assessed the performance of the candidate-selection strategies in two ways. First, we considered how close the predicted F_1_ score from a simulated verification experiment is to that from the overall study. We calculated precision in two modes: ‘default’ (as described in Methods) and ‘weighted’ (where precision scores were modified so that unique calls carried more weight than calls predicted by multiple callers). Second, we assessed the variability in this result across 10 replicate runs of each strategy, allowing us to gauge how much random chance elements of variant-selection perturb the results of a given method (*i.e.* a stability analysis).

Overall, across all simulations, the ‘equal per caller’ approach performs best, showing a negligible mean difference between subset and total F_1_ scores while, additionally, displaying low variability (*i.e.* small spread) in F_1_ score differences across all runs **(Figure 3).** Both the number of algorithms tested and the verification budget size (*i.e.* the number of candidates being selected) factor into which strategy performs optimally. Specifically, when there are large numbers of algorithms or the number of possible verification targets is low, the ‘equal per caller’ method does extremely well (n_targets_ = 100; **Supplementary Figure 1**). By contrast, when the number of verification targets is substantially larger (*i.e.* a considerable proportion of all predictions will be tested), the ‘random rows’ method shows similar performance levels (n_targets_ = 1000 and n_targets_ = 2500; **Supplementary Figures 2 and 3, respectively).** However, the ‘random rows’ method performs poorly when prediction set sizes are highly variable (*i.e.* a small number of callers has a large fraction of the total calls), resulting in some callers with no calls by which to estimate performance. This was the case for runs with verification budgets of n_targets_ = 250 **(Supplementary Figure 4)**, n_targets_ = 500 **(Supplementary Figure 5)** and, in particular, n_targets_ = 100 **(Supplementary Figure 1).** Missing scores were treated as missing data.

**Figure 3:**
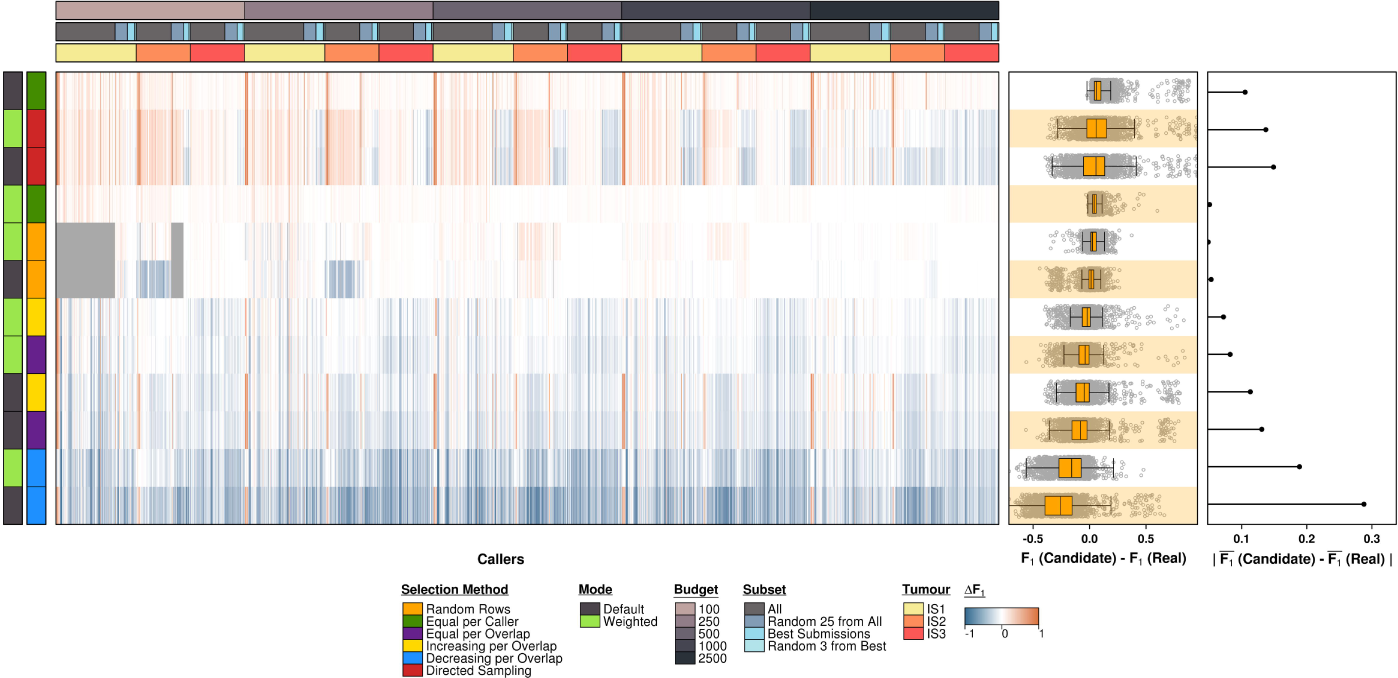
All Simulation Results for Selection Strategy Parameter Combinations. Overall, the best results are obtained using the ‘equal per caller’ method. The ‘random rows’ approach scores comparably except in cases where there is high variability in prediction set sizes across callers. Calls from low-call callers are less likely to be sampled at random and, in cases where none are sampled, it is not possible to get performance estimates for those callers. Failed estimate runs are displayed in grey.

However, the effects of the verification experiment characteristics described above alone do not account for all the variability observed across the simulations. Comparing runs of matching parameter combinations across the three synthetic tumours reveals some inter-tumour differences. Unlike with tumours IS1 **(Supplementary Figure 6)** and IS2 **(Supplementary Figure 7),** the ‘random rows’ method performs best on tumour IS3 suggesting tumour characteristics may have an impact on target selection strategy performance **(Supplementary Figure 8).** The ‘equal per caller’ method is only the second best selection strategy for the IS3 dataset.

We further assessed variability in the results of the selection strategies by running 10 replicate runs of each. The results in **Figure 4** show that the consistency of performance across simulations trends with the overall performance of the selection strategy. An overall positive effect of the adjustment step (‘weighted mode’) on the selection strategies is also visible with the exception of the ‘random rows’ method, on which the weighted precision calculation appears to have no effect. A closer look at the recall and precision scores reveals that the approach with the poorest recall score, ‘decreasing with overlap’ **(Supplementary Figure 9a),** also shows the most sensitivity to the weighted adjustment step in precision calculations **(Supplementary Figure 9b).** Altogether, across methods, recall scores tend to mirror F_1_ scores in both magnitude and amount of spread, which is lower in approaches with higher recall. In contrast, precision scores are highly variable across most selection approaches, regardless of their overall performance.

**Figure 4:**
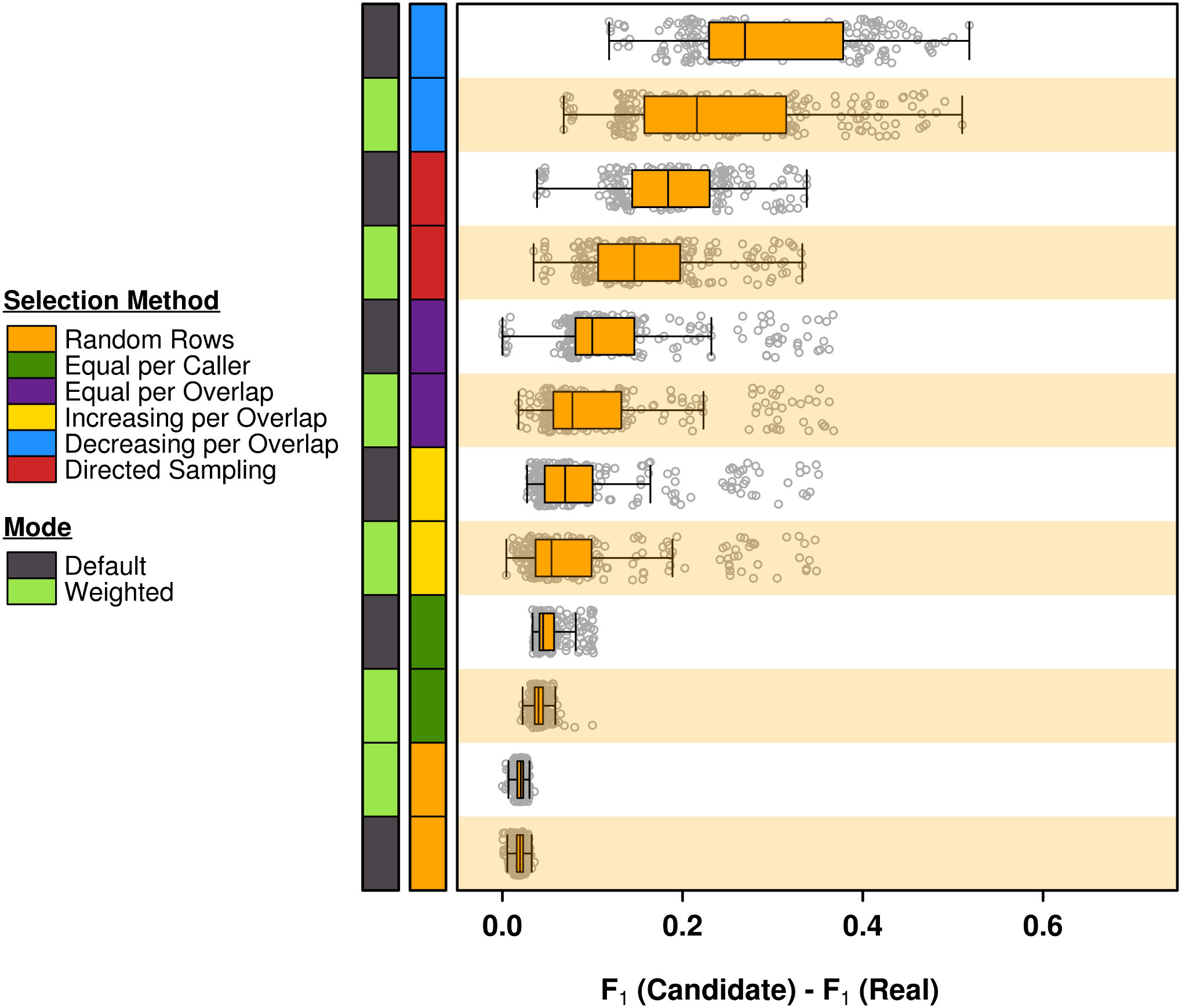
F_1_ Scores Across Replicate Runs. Top selection strategies perform consistently across replicate runs. Strategies are ordered by median scores. The adjustment step in precision calculations improves the ‘equal per caller’ method, but shows little effect on ‘random rows’.

## Discussion

Assessing and comparing the quality of new prediction tools is an important step in their adoption and the truth of their results is arguably the most important component of their quality. When the resources required to independently verify results are substantial, it is vital to choose an unbiased but maximally informative set of results. This is naturally true not just for somatic SNVs, but other predictions like structural variants, fusion proteins, alternative splicing events and epigenetic phenomena, *e.g.* methylation and histone marks. Ongoing research into the error profiles of various data types increases our understanding of what factors influence verification rates [21]. This information helps in distinguishing high- from low-quality calls and goes towards minimizing the amount of prediction verification required. However, with the continuous emergence of new data-generating technologies, *e.g.* third generation sequencing [22], benchmarking studies assessing false positive and false negative rates are likely to remain a fundamental component of computational biological research well into the foreseeable future. Having standardized methods for comparing workflows in contexts such as these will ease the uptake of new techniques more confidently. Valection is a first step towards standardizing and optimizing verification candidate selection.

Evaluation of the target candidate selection approaches presented in this study provides an in-depth view of the effects of call recurrence and algorithm representation on a verification candidate set. Nonetheless, this is by no means an exhaustive set of selection strategies. Although, our findings suggest that the most straightforward approaches (*e.g.* ‘random rows’) are often the most effective, future implementations of more complex strategies may highlight additional factors important to target candidate selection.

The need for informative verification target selections also highlights the importance of simulators for experimental biology, since the best suited method may vary from dataset to dataset. Indeed, as our findings here suggest, optimal candidate-selection strategies for somatic SNV calls may even be affected by various tumour data characteristics. A complete assessment of error profiles is impossible without access to multifarious datasets with an established ground truth. As such, there is a need for reliable simulators in biology to create and analyze gold-standard synthetic datasets to help guide top empirical research. For some time computationally-simulated data has been used to circumvent the difficulties that arise when working with real data [23]. The production of varied synthetic data is comparatively cheap and efficient, restricted only by the computational power and storage space required to generate and hold it. With complete control over data feature profiles, researchers are able to query numerous biological questions simultaneously. As demonstrated here, and specific to cancer genomics, synthetic tumour data can expedite accurate estimation of false negative rates which are difficult to determine in genome-wide mutation calling, thus mitigating the need for large-scale wet lab validation of non-variants. It is important to note, however, that the utility of synthetic data is limited to non-exploratory research. Biological processes or data features that are unknown or poorly understood cannot be adequately simulated, leading to a lack of ‘real-world’ complexity. Therefore, the interplay between experimental and simulated data is critical to the advancement of ‘big data’ disciplines such as genomics. As such, subsequent assessment using comprehensively-characterized real data will be vital to further optimizing candidate-selection strategy.

## Conclusions

Verification of somatic SNV calls made on NGS tumour data is critical due to the high numbers of false positive and false negative calls. However, a thorough search to identify all erroneous calls is a cumbersome and expensive task. Our findings suggest that it may also be an avoidable one. Fewer verification targets may be sufficient to characterize global error rates in data, provided that there is proper optimization of the target candidate selection process. We find that this optimization must factor in not just the scope of the verification study but, conceivably, the characteristics of the dataset itself. To date, few studies have assessed candidate-selection methods for verification purposes. Here, we begin to explore the alternatives available to big data analysts performing confirmatory studies that are both efficient and thorough. By releasing our Valection software publicly, we encourage groups across the wider research community to continue this work. With a straightforward implementation and easy application, Valection has the potential for maximal impact across a wide range of disciplines that rely on verification studies.

## Methods

### Selection Strategies & Software

The **random rows** selection strategy **(Figure 1b)** samples calls at random without replacement from the entire set of calls, and continues until the verification budget has been reached, or there are no more calls left.

The **directed-sampling** selection strategy **(Figure 1c)** begins by constructing a matrix. Row 1 contains all the calls made only by individual callers, row 2 contains the calls made by exactly 2 callers, all the way to row N, which contains the calls that were made by all of the N callers. Each column, j, of the matrix contains only the calls made the j^th^ caller. Note that this means in all rows past 1, calls appear in multiple cells on the same row. Any given cell holds zero or more calls. To select calls, the following procedure is followed for each row, from N to 1, and for each cell in that row, ordered by ascending number of calls:

- Calculate the cell budget as the total remaining verification budget divided among the yet unexamined cells in the rest of the matrix.
- Select calls without replacement from the cell in question up to the cell budget (these calls become invalid selections for future cells). Each call selected reduces the total remaining verification budget.
- If any budget remains once all cells have been selected from, the process is repeated.

The **equal per caller** selection strategy **(Figure 1d)** divides the verification budget equally among all callers. The set of calls that each individual caller made is sampled from without replacement up to that caller’s portion of the total budget. A call selected by one caller becomes an invalid choice for all other callers. If a single caller does not have enough available calls (calls not yet selected in another caller’s budget), its remaining budget is distributed equally to the other callers.

The **equal per overlap** selection strategy **(Figure 1e)** is based around the number of times each call was made. With N callers, the verification budget is divided N ways. Out of the set of calls made only once (all the calls unique to any caller), calls are selected without replacement up to the sub-budget. This is repeated for all the calls made by exactly two callers, and so on up every level of overlap. If a single level of overlap does not have enough available calls (calls not yet selected in another overlap level’s budget), its remaining budget is distributed equally to the other levels.

The **increasing with overlap** selection strategy **(Figure 1f)** is similar to equal per overlap, but instead of selecting an equal number of calls at every level of overlap, it selects a number from each level of overlap proportional to the level of overlap.

The **decreasing with overlap** selection strategy **(Figure 1g)** is identical to increasing with overlap, but the number of calls selected at each level is inversely proportional to the level of overlap.

All of these methods are available through four commonly used programming languages C, Perl, Python and R. The implementations have robust user-level documentation and are openly available at both their appropriate public repositories (*i.e.* CPAN, PyPI and CRAN) and on our website at: labs.oicr.on.ca/boutros-lab/software/valection.

The selection strategy algorithms were implemented in C, and compiled using the GNU Compiler Collection (v4.8.1). The implementations also made use of GLib (v 2.44.0). The R statistical environment (v3.1.3) was used for statistical analysis and data subsetting. Perl (v5.18.2) was used to coordinate the simulations. All plots were generated with the same version of R using the “BPG” (v5.2.8) [24], “lattice” (v0.20-31) and “latticeExtra” (v0.6-26) packages. The analysis scripts are also available at http://labs.oicr.on.ca/boutros-lab/software/valection.

### Simulated Data

To test the accuracy of these different approaches empirically, we applied them to gold-standard data from the ICGC-TCGA DREAM Somatic Mutation Calling Challenge [20]. This is a global crowd-sourced benchmarking competition aiming to define the optimal methods for the detection of somatic mutations from NGS-based whole-genome sequencing. The challenge has two components, one using simulated data created using BAMSurgeon software [4] and the other using experimentally-verified analyses of primary tumours. To test the accuracy of our approaches on representation algorithms, we exploited the SNV data from the first three *in silico* tumours. This dataset comprises 261 genome-wide prediction sets made by 21 teams and there are no access restrictions. The raw BAM files are available at SRA with IDs SRX570726, SRX1025978 and SRX1026041. Truth files are available as VCFs at https://www.synapse.org/#!Synapse:syn2177211. Prediction-by-submission matrices for all submissions are provided in Supplementary Tables 1-3, as well as the best submissions from each team in Supplementary Table 4, truth calls in Supplementary Tables 5-7 and a confusion matrix in Supplementary Table 8.

To probe a range of possible verification studies, we ran a very broad set of simulations. For each run, we pre-specified a tumour, a number of algorithms and a number of mutations to be selected for verification, and ran each of the candidate-selection strategies listed above. We then calculated the F_1_ score (along with precision and recall) based on the verification study, assuming verification results are ground truth. Finally, we compared the true F_1_ for a given algorithm on a given tumour across all mutations to the one inferred from the verification experiment.

We used three separate tumours with diverse characteristics (https://www.synapse.org/#!Synapse:syn312572/wiki/62018), including a range of tumour cellularities and the presence or absence of sub-clonal populations. We selected subsets of algorithms for benchmarking in four different ways:

i. the complete dataset (X)
ii. the single best submission from each team (X-best)
iii. three randomly selected entries from X-best (repeated 10 times)
iv. 25 randomly selected entries from X (repeated 10 times)

Lastly, we considered verification experiment sizes of 100, 250, 500, 1000 and 2500 candidates per tumour. Thus, in total, we analyzed each of the candidate-selection algorithms in 22 datasets for 3 tumours and 5 verification sizes, for 330 total comparisons.

### Statistical Analyses

The precision, recall and F_1_ score of each caller were calculated as follows, from the caller’s true positive (TP), false positive (FP) and false negative (FN) values, as estimated by the selection strategy. Here, FNs are true calls sampled by the selection strategy that were not made by the caller in question (*i.e.* another caller made it).

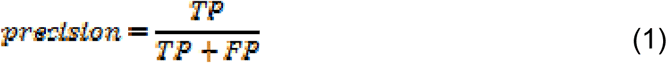

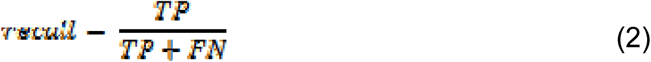

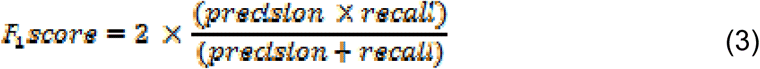

When no calls were selected to calculate a value for a caller, scores were given values of N/A. This happened primarily with the ‘random rows’ method.

Additionally, each precision score was calculated in an adjusted and unadjusted manner. A caller’s precision in the unadjusted form was calculated exactly as described above, using all the calls made by the caller and selected for verification as the TPs and FPs. In the adjusted form, the selected calls were first divided into groups, according to how many callers made the call. Then, the precision was calculated separately using the calls from each group. The final precision was calculated as a weighted average of the precision of each group of calls, with weights equal to the total number of calls (verified and unverified) that caller made at that overlap level. Thus, in a two-caller example, a caller that made 100 unique calls and 50 calls shared with the other caller would count its precision from unique calls twice as strongly as its precision from shared calls.

## List of abbreviations

SNV: single-nucleotide variant
NGS: next-generation sequencing
ICGC: International Cancer Genome Consortium
TCGA: The Cancer Genome Atlas
DREAM: Dialogue for Reverse Engineering Assessments and Methods
SMC-DNA: Somatic Mutation Calling DNA Challenge
TP: true positive
FP: false positive
FN: false negative

## Declarations

### Ethics approval and consent to participate

Not applicable

### Consent for publication

Not applicable

### Availability of data and material

The datasets supporting the conclusions of this article are included in its additional files and in the Supplementary of Ewing *et al.* [4]. The main Valection project page is at:

http://labs.oicr.on.ca/boutros-lab/software/valection

Programmatic bindings for the source-code are additionally available at: https://pypi.python.org/pypi/valection/1.0.1 http://search.cpan.org/dist/Bio-Sampling-Valection/

### Authors’ contributions

Initiated the project: CIC, JMS, PCB

Data preparation: TNY, CC, KEH

Generated tools and reagents: CIC, DY, TNY, CP, KE

Performed statistical and bioinformatics analyses: CIC, DY, DHS, TNY, KEH

Supervised research: MF, KE, AAM, RGB, JMS, PCB

Wrote the first draft of the manuscript: DHS, PCB

Approved the manuscript: all authors

## Acknowledgements

The authors thank Dr. Jared Simpson and Dr. John McPherson for thoughtful discussions and all members of the Boutros lab and the SMC-DNA team for helpful suggestions.

## Additional files

**Additional file 1: Supplementary Figure 1.**

TIFF 9.4 Mb

Simulations with 100 verification targets, across all tumours. The ‘equal per caller’ method (weighted mode) performs optimally as the ‘random rows’ method generates N/As.

**Additional file 2: Supplementary Figure 2.**

TIFF 9.2 Mb

All simulations with 1000 verification targets, across all tumours. The best results come from the ‘random rows’ and the ‘equal per caller’ (weighted mode) methods.

**Additional file 3: Supplementary Figure 3.**

TIFF 8.9 Mb

All simulations with 2500 verification targets, across all tumours. The best results come from the ‘random rows’ and the ‘equal per caller’ (weighted mode) methods.

**Additional file 4: Supplementary Figure 4.**

TIFF 9.9 Mb

All simulations with 250 verification targets, across all tumours. The ‘equal per caller’ method (weighted mode) performs optimally as the ‘random rows’ method generates N/As.

**Additional file 5: Supplementary Figure 5.**

TIFF 9.6 Mb

All simulations with 500 verification targets, across all tumours. The ‘equal per caller’ method (weighted mode) performs optimally as the ‘random rows’ method generates N/As.

**Additional file 6: Supplementary Figure 6.**

TIFF 18 Mb

All simulations for tumour IS1. Optimal results are achieved with the ‘equal per caller’ method (weighted mode).

**Additional file 7: Supplementary Figure 7.**

TIFF 12 Mb

All simulations for tumour IS2. Optimal results are achieved with the ‘equal per caller’, ‘increasing per overlap’ and ‘equal per overlap’ methods (weighted mode).

**Additional file 8: Supplementary Figure 8.**

TIFF 14 Mb

All simulations for tumour IS3. Optimal results are achieved with the ‘random rows’ method, regardless of how precision is calculated.

**Additional file 9: Supplementary Figure 9.**

TIFF 4.1 Mb

a) Recall scores from all runs, displayed per candidate-selection strategy. b) Precision scores from all runs, calculated with and without a weight adjustment step (default mode and weighted mode, respectively) and displayed per candidate-selection strategy.

**Additional file 10: Supplementary Table 1.**

CSV 57 Mb

A prediction-by-submission matrix of all SNV call submissions for tumour IS1 where SNV predictions are annotated with chromosome (“CHROM”) and position (“END”).

**Additional file 11: Supplementary Table 2.**

CSV 29 Mb

A prediction-by-submission matrix of all SNV call submissions for tumour IS2 where SNV predictions are annotated with chromosome (“CHROM”) and position (“END”).

**Additional file 12: Supplementary Table 3.**

CSV 3.6 Mb

A prediction-by-submission matrix of all SNV call submissions for tumour IS3 where SNV predictions are annotated with chromosome (“CHROM”) and position (“END”).

**Additional file 13: Supplementary Table 4.**

CSV 3.3 kb

A summary table of the top team submissions for each tumour, includes submission ID, team alias, the number of true positives, true negatives, false positives and false negatives, as well as the precision, recall and F_1_ scores.

**Additional file 14: Supplementary Table 5.**

CSV 3.1 Mb

A table of all predicted SNVs for tumour IS1, annotated by chromosome (“chrom”) and position (“pos”), and a “truth” column for whether the call is a true positive (1) or not (0).

**Additional file 15: Supplementary Table 6.**

CSV 2.5 Mb

A table of all predicted SNVs for tumour IS2, annotated by chromosome (“chrom”) and position (“pos”), and a “truth” column for whether the call is a true positive (1) or not (0).

**Additional file 16: Supplementary Table 7.**

CSV 329 kb

A table of all predicted SNVs for tumour IS3, annotated by chromosome (“chrom”) and position (“pos”), and a “truth” column for whether the call is a true positive (1) or not (0).

**Additional file 17: Supplementary Table 8.**

CSV 20 kb

A summary table of all submissions from across all tumours, includes submission ID, the number of true positives, true negatives, false positives and false negatives, as well as the precision, recall and F_1_ scores.

## References

1. Abnikova I, Leonard S, Skelly T, Brown A, Jackson D, Gourtovaia M, Qi G, Te Boekhorst R, Faruque N, Lewis K, Cox T. Analysis of context-dependent errors for Illumina sequencing. J Bioinform Comput Biol. 2012 Apr;10(2):1241005.

2. Meacham F, Boffelli D, Dhahbi J, Martin DIK, Singer M, Pachter L. Identification and correction of systemic error in high-throughput sequence data. BMC Bioinformatics. 2011 Nov 21;12:4511.

3. Derrien T, Estellé J, Marco Sola S, Knowles DG, Raineri E, Guigo R, Ribeca P. Fast computation and applications of genome mappability. PLoS One. 2012;7(1):e30377.

4. Ewing AD, Houlahan KE, Hu Y, Ellrott K, Caloian C, Yamaguchi TN, Bare JC, P’ng C, Waggott D, Sabelnykova VY, ICGC-TCGA DREAM Somatic Mutation Calling Challenge participants, Kellen MR, Norman TC, Haussler D, Friend SH, Stolovitzky G, Margolin AA, Stuart JM, Boutros PC. Combining tumor genome simulation with crowdsourcing to benchmark somatic single-nucleotide-variant detection. Nat Methods. 2015 Jul;12(7):623-30.

5. Hofmann AL, Behr J, Singer J, Kuipers J, Beisel C, Schraml P, Moch H, Beerenwinkel N. Detailed simulation of cancer exome sequencing data reveals differences common limitations of variant callers. BMC Bioinformatics. 2017 Jan 3;18(1):8. doi: 10.1186/s12859-016-1417-7.

6. Lam HYK, Clark MJ, Chen R, Chen R, Natsoulis G, O’Huallachain M, Dewey FE, Habegger L, Ashley EA, Gerstein MB, Butte AJ, Ji HP, Snyder M. Performance comparison of whole-genome sequencing platforms. Nat Biotechnol. 2012 Jan;30(1):78-82.

7. Wall JD, Tang LF, Zerbe B, Kvale MN, Kwok P-Y, Schaefer C, Risch N. Estimating genotype error rates from high-coverage next-generation sequence data. Genome Res. 2014 Nov;24(11):1734-9.

8. Quon G, Haider S, Deshwar AG, Cui A, Boutros PC, Morris Q. Computational purification of individual tumor gene expression profiles leads to significant improvements in prognostic prediction. Genome Med. 2013 Mar 28;5(3):29.

9. Carter SL, Cibulskis K, Helman E, McKenna A, Shen H, Zack T, Laird PW, Onofrio RC, Winckler W, Weir BA, Beroukhim R, Pellman D, Levine DA, Lander ES, Meyerson M, Getz G. Absolute quantification of somatic DNA alterations in human cancer. Nat Biotechnol. 2012 May;30(5):413-21.

10. Roth A, Khattra J, Yap D, Wan A, Laks E, Biele J, Ha G, Aparicio S, Bouchard-Cote A, Shah SP. PyClone: statistical inference of clonal population structure in cancer. Nat Methods. 2014 Apr;11(4):396-8.

11. Lee LG, Connell CR, Woo SL, Cheng RD, McArdle BF, Fuller CW, Halloran ND, Wilson RK. DNA sequencing with dye-labeled terminators and T7 DNA polymerase: effect of dyes and dNTPs on incorporation of dye-terminators and probability analysis of termination fragments. Nucleic Acids Res. 1992 May 25;20(10):2471-83.

12. Rehm HL, Bale SJ, Bayrak-Toydemir P, Berg JS, Brown KK, Deignan JL, Friez MJ, Funke BH, Hegde MR, Lyon E, Working Group of the American College of Medical Genetics and Genomics Laboratory Quality Assurance Commitee. ACMG clinical laboratory standards for next-generation sequencing. Genet Med. 2013 Sep;15(9):733-47.

13. Sikkema-Raddatz B, Johansson LF, de Boer EN, Almomani R, Boven LG, van den Berg MP, van Spaendonck-Zwarts KY, van Tintelen JP, Sijmons RH, Jongbloed JD, Sinke RJ. Targeted next-generation sequencing can replace Sanger in clinical diagnostics. Hum Mutat. 2013 Jul;34(7):1035-42.

14. Nelson AC, Bower M, Baughn LB, Henzler C,Onsongo G, Silverstein KAT, Schomaker M, Deshpande A, Beckman KB, Yohe S, Thyagarajan B. Criteria for clinical reporting of variants from a broad target capture NGS assay without Sanger verification. JSM Biomark. 2015;2(1):1005.

15. Strom SP, Lee H, Das K, Vilain E, Nelson SF, Grody WW, Deignan JL. Assessing the necessity of confirmatory testing for exome-sequencing results in a clinical molecular diagnostic laboratory. Genet Med. 2014 Jul;16(7):510-5.

16. Cancer Genome Atlas Network. Comprehensive genomic characterization of head and neck squamous cell carcinomas. Nature. 2015 Jan 29;517(7536):576-82.

17. Chong LC, Albuquerque MA, Harding NJ, Caloian C, Chan-Seng-Yue M, de Borja R, Fraser M, Denroche RE, Beck TA, van der Kwast T, Bristow RG, McPherson JD, Boutros PC. SeqControl: process control for DNA sequencing. Nat Methods. 2014 Oct; 11 (10): 1071-5.

18. Lee W, Jiang Z, Liu J, Haverty PM, Guan Y, Stinson J, Yue P, Zhang Y, Pant KP, Bhatt D, Ha C, Johnson S, Kennemer MI, Mohan S, Nazarenko I, Watanabe C, Sparks AB, Shames DS, Gentleman R, de Sauvage FJ, Stern H, Pandita A, Ballinger DG, Drmanac R, Modrusan Z, Seshagiri S, Zhang Z. The mutation spectrum revealed by paired genome sequences from a lung cancer patient. Nature. 2010 May 27;465(7297):473-7.

19. Ratan A, Miller W, Guillory J, Stinson J, Seshagiri S, Schuster SC. Comparison of sequencing platforms for single nucleotide variant calls in a human sample. PLoS One. 2013 Feb 6;8(2):e55089. doi: 10.1371/journal.pone.0055089.

20. Boutros PC, Margolin AA, Stuart JM, Califano A, Stolovitzky G. Toward better benchmarking: challenge-based methods assessment in cancer genomics. Genome Biol. 2014 Sep 17;15(9):462. doi: 10.1186/s13059014-0462-7.

21. Park M-H, Rhee H, Park JH, Woo HM, Choi BO, Kim BY, Chung KW, Cho YB, Kim HJ, Jung JW, Koo SK. Comprehensive analysis to improve the validation rate for single nucleotide variants detected by next-generation sequencing. PLoS One. 2014 Jan 29;9(1):e86664.

22. Lee H, Gurtowski J, Yoo S, Nattestad M, Marcus S, Goodwin S, McCombie WR, Schatz M. Third-generation sequencing and the future of genomes. bioRxiv 048603; doi:https://doi.org/10.1101/048603.

23. Escalona M, Rocha S, Posada D. A comparison of tools for the simulation of genomic next-generation sequencing data. Nat Rev Genet. 2016 Aug;17(8):459-69.

24. P’ng C, Green J, Chong LC, Waggott D, Prokopec SD, Shamsi M, Nguyen F, Mak DYF, Lam F, Albuquerque MA, Wu Y, Jung EH, Starmans MHW, Chan-Seng-Yue MA, Yao CQ, Liang B, Lalonde E, Haider S, Simone NA, Sendorek D, Chu KC, Moon NC, Fox NS, Grzadkowski MR, Harding NJ, Fung C, Murdoch AR, Houlahan KE, Wang J, Garcia DR, de Borja R, Sun RX, Lin X, Chen GM, Lu A, Shiah Y-J, Zia A, Kearns R, Boutros P. BPG: seamless, automated and interactive visualization of scientific data. bioRxiv 156067; doi:https://doi.org/10.1101/156067.

